# Genomic alterations in persistently infecting oncolytic Newcastle disease virus reveal mechanisms of viral persistence in bladder cancer cells

**DOI:** 10.1101/2025.05.06.652422

**Authors:** U Ahmad, D Chau, SC Chan, K Yusoff, A Veerakumarasivam, S Abdullah

## Abstract

Newcastle disease virus (NDV) is a promising oncolytic agent with a non-segmented, negative-sense single-stranded RNA (ssRNA) genome of approximately 15 kb. While NDV selectively replicates in and lyses a wide range of human cancer cells, a subset of these cells develop persistent infections, potentially compromising the therapeutic efficacy of NDV-based treatments. To investigate the molecular basis of persistent infection, we performed transcriptome profiling of TCCSUP bladder cancer cells persistently infected with the NDV AF2240 strain. Deep sequencing using Illumina HiSeq 2000 was conducted in triplicate, and the resulting viral and host reads were separated and analyzed. Using Integrative Genomic Viewer (IGV) software, we identified several nucleotide variants linked to persistence. Specifically, nucleotide alterations included a deletion at 359A→C and a substitution at 1653C→T within the nucleoprotein (NP) gene, as well as an insertion at 3338C→T in the matrix protein (M) gene. Additionally, a GGG base insertion was detected at position 2290 in the phosphoprotein (P) gene. Crucially, we observed truncations in the hemagglutinin-neuraminidase (HN) gene (nt 8263–8390) and the large polymerase (L) gene (nt 6203–6342). These mutations and truncations suggest significant disruptions in viral replication, assembly, and host cell attachment, potentially facilitating viral persistence in bladder cancer cells. Understanding these genomic alterations provides valuable insights into the mechanisms driving viral persistence and could inform strategies to optimize the oncolytic efficacy of NDV in cancer therapy.

## 1. Introduction

Newcastle disease virus (NDV), a member of the *Avian paramyxovirus* (APMV) family, has demonstrated significant oncolytic potential against various cancer types. NDV contains a non-segmented, negative-sense single-stranded RNA (ssRNA) genome of approximately 15– 16 kb in size, which encodes six structural proteins arranged as 3’-NP-P-M-F-HN-L-5’[1, 2]. These structural proteins include the nucleoprotein (NP, 55 kDa), phosphoprotein (P, 53 kDa), and large polymerase protein (L, 200 kDa), which form the nucleocapsid[3, 4]. The external membrane of the virion is composed of the haemagglutinin-neuraminidase (HN, 74 kDa) and fusion glycoprotein (F, 67 kDa), crucial for viral attachment and fusion with host cells[5, 6]. The inner layer of the viral envelope is made of the matrix protein (M, 40 kDa), which plays an essential role in viral assembly and budding[7].

NDV’s ability to selectively replicate in and kill cancer cells has been well-established in both preclinical and clinical studies[8, 9]. However, it has been observed that a subset of cancer cells can develop persistent infections, enabling them to resist NDV-mediated oncolysis[10]. This phenomenon of viral persistence has also been noted in other oncolytic viruses, such as Reovirus[11] and the Edmonston Measles virus[12]. Persistent infection suggests that a subpopulation of cancer cells may harbor genetic or molecular mechanisms that allow them to evade viral lysis[10]. This poses a significant challenge to the therapeutic efficacy of NDV, as tumors contain genetically heterogeneous cell populations with various aberrations[13].

In heterogeneous tumors, the differential response to NDV could be attributed to the diverse genetic profiles of the cancer cells[14]. Persistent infection is particularly concerning as it may reduce the overall effectiveness of NDV-based oncolytic virotherapy[15]. Although recombinant NDV strains with enhanced oncolytic potential have been developed[16], the risk of viral persistence remains. Understanding the molecular mechanisms that allow cancer cells to resist NDV-induced cytotoxicity is essential for improving therapeutic outcomes.

In this study, we employed bioinformatics tools to investigate the genomic profile of the persistently infecting NDV strain (NDVpi) isolated from TCCSUP bladder cancer cells. Using RNA-Seq data, we identified several nucleotide variations, including deletions, insertions, and truncations in key viral proteins. These mutations likely contribute to viral persistence in cancer cells. By deciphering these genomic alterations, we aim to gain insights into the mechanisms driving viral persistence and suggest strategies to enhance NDV’s oncolytic efficacy.

## 2. Materials and Methods

### 2.1. Cell Culture and Virus Infection

The TCCSUP human bladder cancer cell line was used as the in vitro model for persistent infection studies. Cells were maintained in RPMI-1640 medium supplemented with 10% fetal bovine serum (FBS), 1% penicillin-streptomycin, and 1% L-glutamine, under standard conditions at 37°C with 5% CO □. The cells were persistently infected with the oncolytic *Newcastle disease virus* (NDV) AF2240 strain at a multiplicity of infection (MOI) optimized for persistent infection studies[10]. Infections were monitored over time to ensure establishment of persistent infection without full cytolysis. At the point of harvest, cells were washed with PBS, lysed, and total RNA was extracted for transcriptomic analysis.

### 2.2. RNA Extraction and Quality Control

Total RNA was extracted from both the persistently infected TCCSUP (TCCSUPPi) cells and uninfected TCCSUP control cells using the Qiagen RNeasy Micro Kit (Qiagen, Germany), following the manufacturer’s protocol. RNA concentration and purity were measured using a NanoDrop spectrophotometer (Thermo Fisher Scientific), while RNA integrity was assessed using an Agilent 2100 Bioanalyzer (Agilent Technologies, Santa Clara, CA, USA). Only RNA samples with an RNA Integrity Number (RIN) greater than 7.0 were considered suitable for sequencing to ensure high-quality transcriptomic data.

### 2.3. RNA Sequencing

RNA-Seq libraries were prepared from 1 µg of total RNA using the TruSeq RNA Library Preparation Kit (Illumina) following the manufacturer’s protocol. Briefly, mRNA was isolated from total RNA using poly(A) selection, followed by fragmentation and reverse transcription into cDNA. The cDNA was then end-repaired, A-tailed, and ligated with Illumina adapters. The libraries were amplified by PCR and purified using AMPure XP beads (Beckman Coulter). Library quality was validated using the Agilent 2100 Bioanalyzer, and libraries were quantified using qPCR.

The prepared libraries were sequenced on an Illumina HiSeq 2000 platform, generating paired-end reads of 100 bp in triplicate for each sample. Approximately 30 million reads were generated per sample to ensure sufficient coverage for both host and viral transcriptome analysis.

### 2.4. Bioinformatics Analysis

Raw sequencing reads were first subjected to quality control using FastQC[17]. Low-quality reads and adapter sequences were trimmed using Trimmomatic. The high-quality filtered reads were then mapped to a combined human (hg19) and NDV reference genome (GenBank: JX012096.20) using HISAT2 aligner[18]. After mapping, BAM files for viral and human reads were separated for downstream analysis.

### 2.5. Viral Genome Analysis

The viral BAM file was analyzed using the Integrative Genomic Viewer (IGV)[19] to visualize aligned reads and identify mutations in the NDV genome. To facilitate this, a custom viral genome annotation file was generated using the NDV reference genome and corresponding GFF3 and FASTA files. Viral variants, including single nucleotide polymorphisms (SNPs), insertions, deletions, and truncations, were manually identified and recorded. Supplementary workflow details are provided in Supplementary Figure 1.

## 3. Results and Discussion

### Genomic Variations in Persistently Infecting NDV (NDVpi)

To investigate the genomic alterations in the Newcastle disease virus (NDV) strain AF2240 responsible for persistent infection in bladder cancer cells, RNA-Seq data from TCCSUPPi cells were analyzed. The viral genome was aligned to the NDV reference sequence using the Integrative Genomic Viewer (IGV) to identify mutations, insertions, deletions, and truncations. Throughout the viral genome (3’-NP-P-M-F-HN-L-5’), spanning regions revealed several nucleotide mismatches and low-quality bases (often semi-transparent or faint), which were detected at key coding regions in the viral genome (Table 1). These alterations suggest mutations that may contribute to the establishment and maintenance of persistent infection by the virus.

**Table 1:**
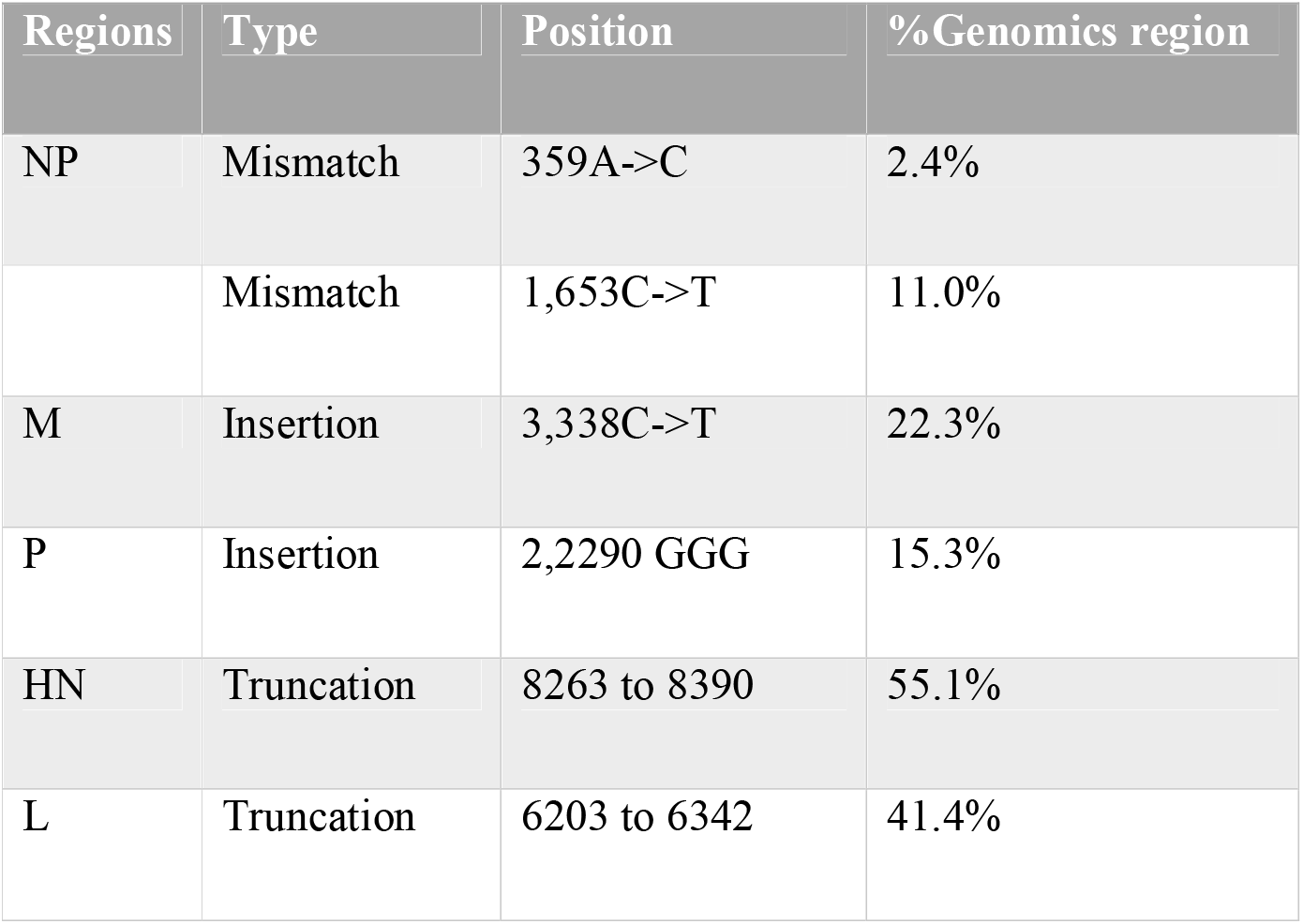
Putative variants detection

| Regions | Type | Position | % Genomics region |
| --- | --- | --- | --- |
| NP | Mismatch | 359A->C | 2.4% |
|  | Mismatch | 1,653C->T | 11.0% |
| M | Insertion | 3,338C->T | 22.3% |
| P | Insertion | 2,2290 GGG | 15.3% |
| HN | Truncation | 8263 to 8390 | 55.1% |
| L | Truncation | 6203 to 6342 | 41.4% |

#### 1. Nucleoprotein (NP) Variants

The nucleoprotein (NP) gene displayed two distinct mismatches, including a deletion at position 359A→C (2.4% of the reads) and a substitution at position 1653C→T (11.0% of the reads). NP plays a critical role in encapsidating viral RNA, a process essential for viral replication and transcription[1]. Mutations in this region may impact the stability of the ribonucleoprotein complex, potentially impairing viral replication and allowing a subset of infected cells to survive, thereby contributing to viral persistence (Figure 2). In other studies of RNA viruses, similar NP mutations have been linked to altered replication efficiency and immune evasion[9].

**Figure 1.**
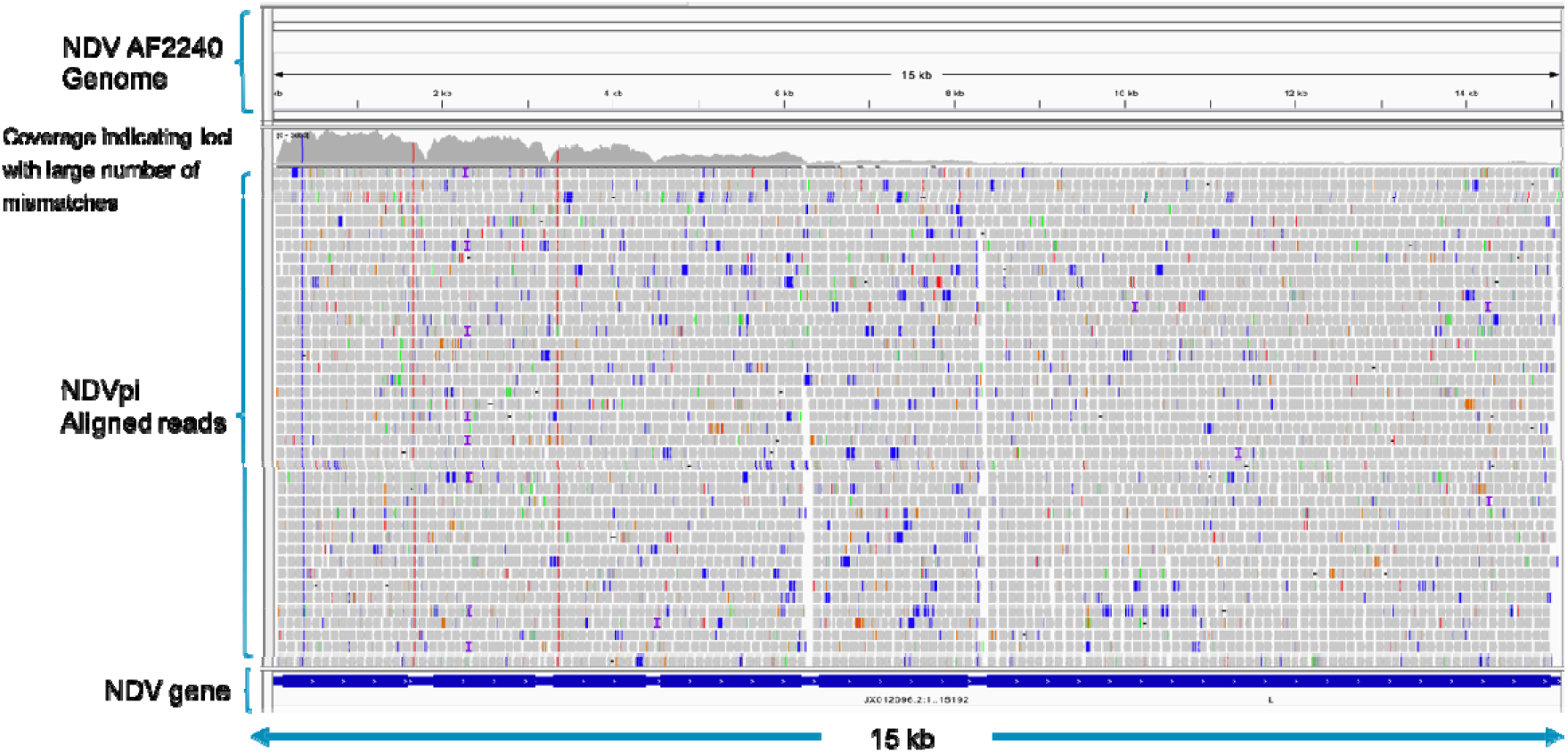
IGV alignment of deep sequencing reads of NDVpi obtained from cancer RNA-Seq data. The various proteins encoded by the corresponding regions of the genome are shown above the alignment. The horizontal bars represent read alignments with the genome; gray bars are properly aligned and placed read pairs, and coloured bars are read pairs that differ from the expected insert size or read orientation. The vertical coloured lines in the coverage plot represent nucleotide positions that were different from the reference genome sequence.

**Figure 2.**
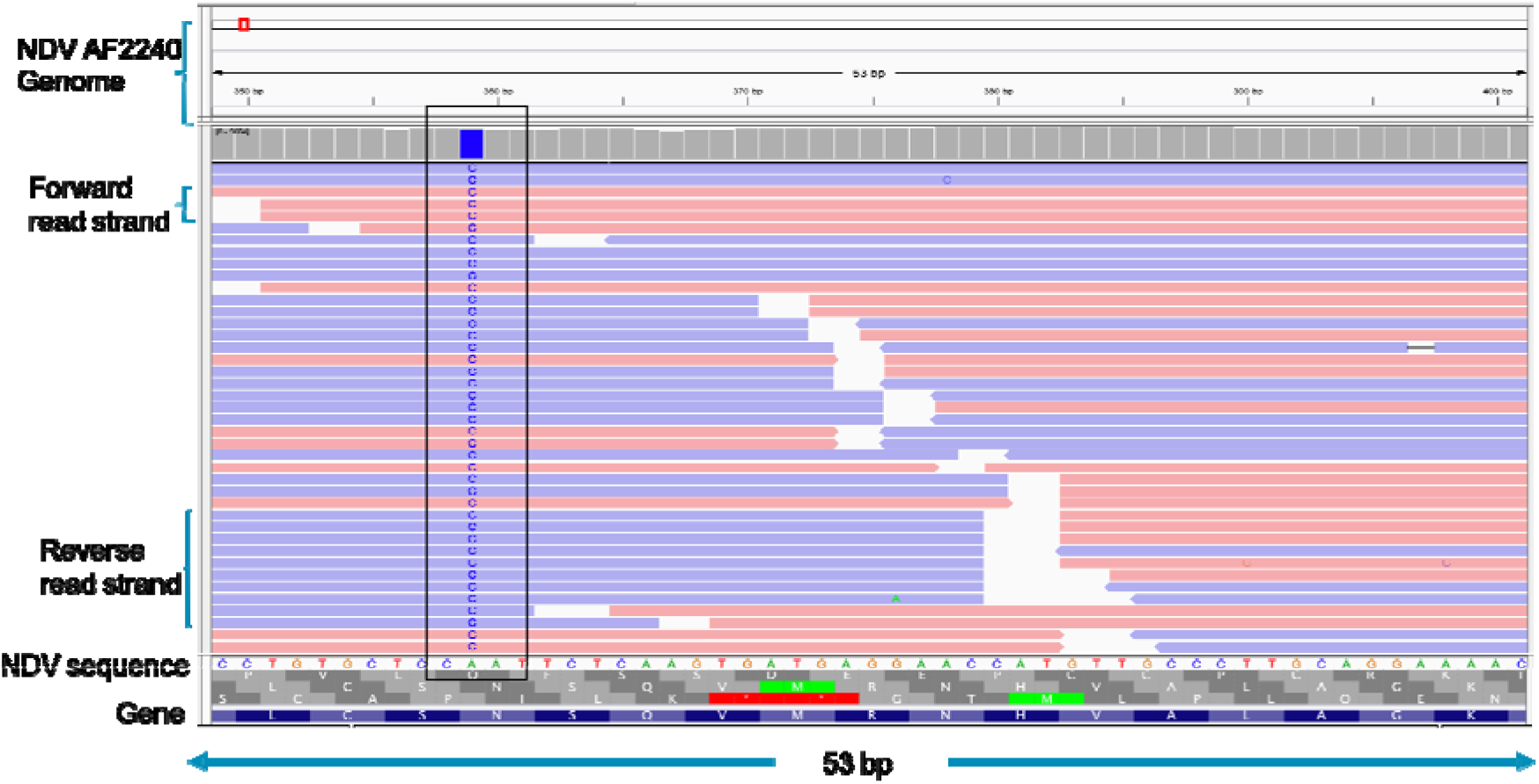
Putative variant in NP protein of NDVpi. Changes in nucleotide sequence of NP protein is observed at 359AdelC (deletion) and 1,653C>T positions.

#### 2. Matrix Protein (M) Variants

A single insertion at 3338C→T was identified in the matrix protein (M) gene, accounting for 22.3% of the viral reads (Table 1). The M protein is involved in viral assembly and budding from the host cell membrane[7]. Mutations in this protein could reduce the efficiency of virion assembly or release, limiting cytopathic effects and allowing infected cells to remain viable, which can facilitate persistent infection (Figure 3). Previous research has shown that alterations in matrix proteins can significantly affect viral budding efficiency, promoting persistent infections[8].

**Figure 3.**
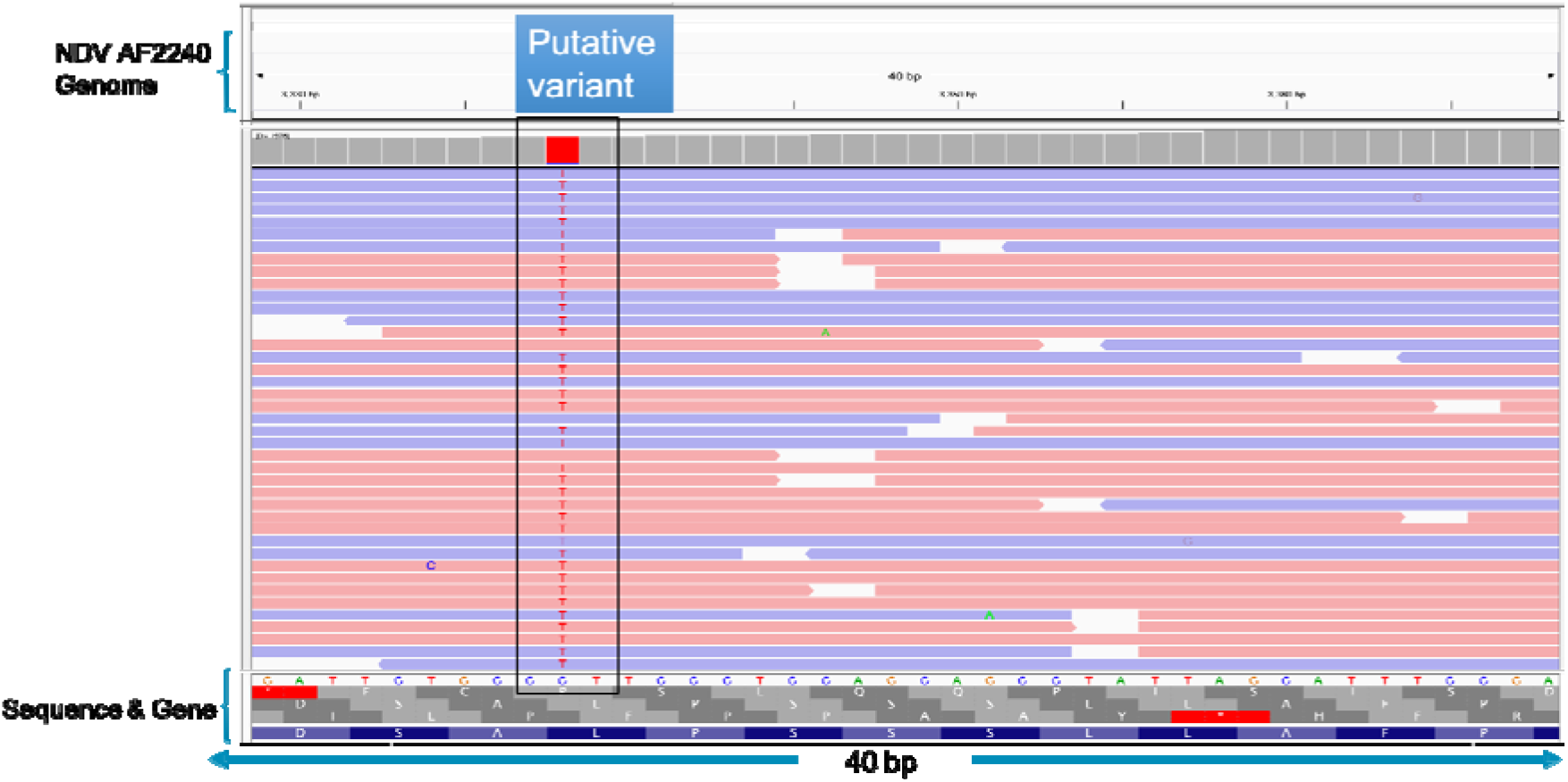
Putative variant in M protein of NDVpi. Changes in nucleotide sequence of M protein is observed at 3,338CinsT (insertion).

#### 3. Phosphoprotein (P) Variants

An insertion of three guanine nucleotides (GGG) was detected at position 2290 in the phosphoprotein (P) gene, contributing to 15.3% of the aligned reads. The P protein is essential for the viral RNA-dependent RNA polymerase complex, working as a cofactor in replication and transcription[11]. Mutations in this region could affect the functionality of the viral polymerase, leading to inefficient viral RNA synthesis, which can contribute to viral persistence through suboptimal replication dynamics (Table 1). Changes in polymerase cofactor proteins are known to be crucial in maintaining persistent infections by modulating viral replication and host immune response[5].

#### 4. Hemagglutinin-Neuraminidase (HN) and Large Polymerase (L) Truncations

The IGV analysis also revealed significant truncations in the hemagglutinin-neuraminidase (HN) and large polymerase (L) genes. A truncation between nucleotide positions 8263 and 8390 in the HN gene was detected in 55.1% of the viral reads (Table 1). The HN protein is vital for viral attachment and entry into host cells by binding to sialic acid receptors on the cell surface[16]. Truncations in this region may impair the virus’s ability to efficiently infect new host cells, limiting its spread and contributing to viral persistence (Figure 4). Similarly, the L gene, which encodes the viral RNA polymerase, exhibited a truncation between nucleotide positions 6203 and 6342 in 41.4% of the reads. Truncations in the L gene can significantly reduce viral replication capacity[10, 20]. Such genome truncations have been observed in other RNA viruses as well, where they contribute to persistent infection by limiting the cytolytic potential of the virus[12].

**Figure 4.**
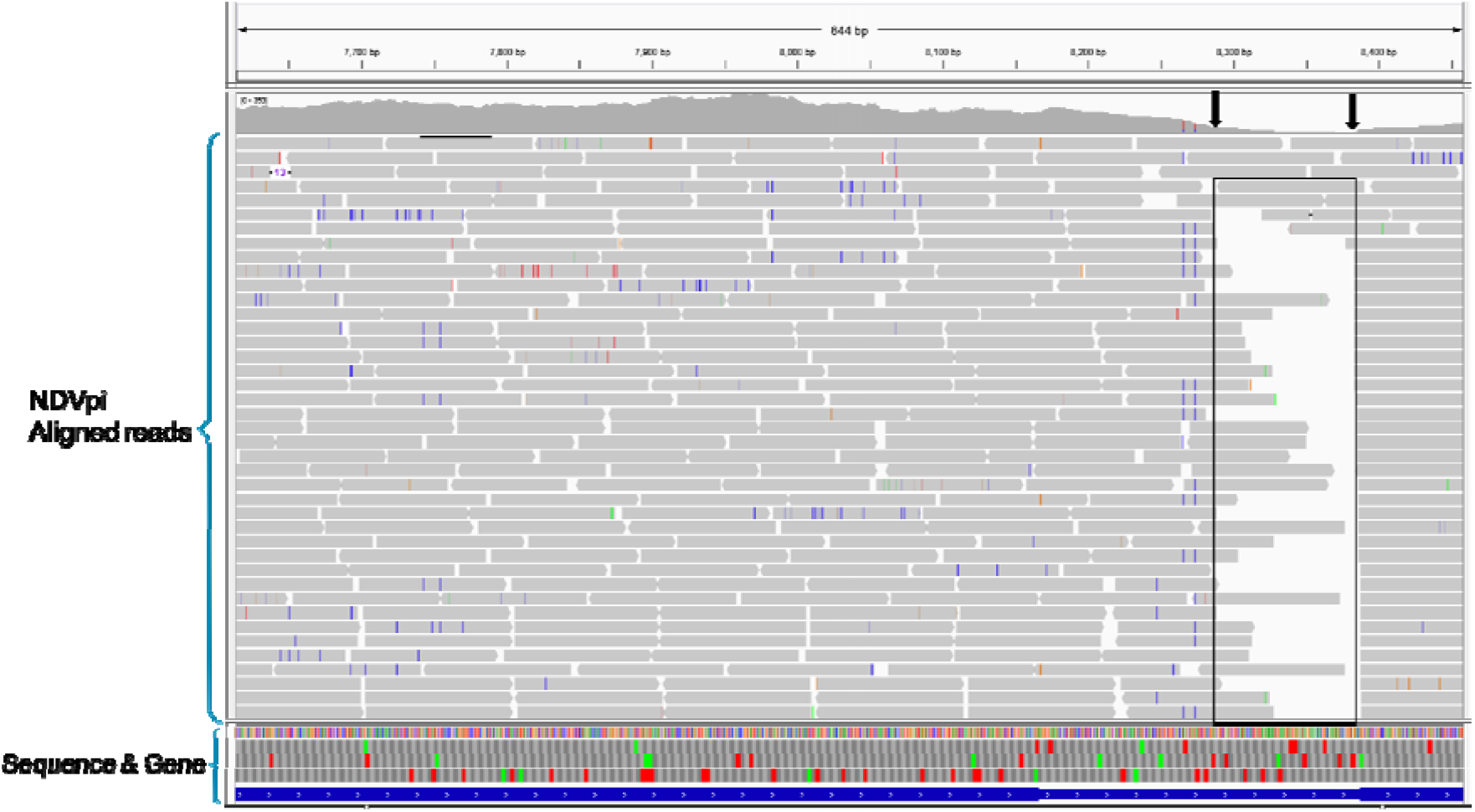
NDVpi aligned to the reference sequence showing genome truncation. Alignment of deep-sequence reads obtained from NDVpi. The alignment shows genome truncations (indicated by arrows), the majority of which were mapped to nucleotide position from 8263 to 8390 in HN protein.

**Figure 5.**
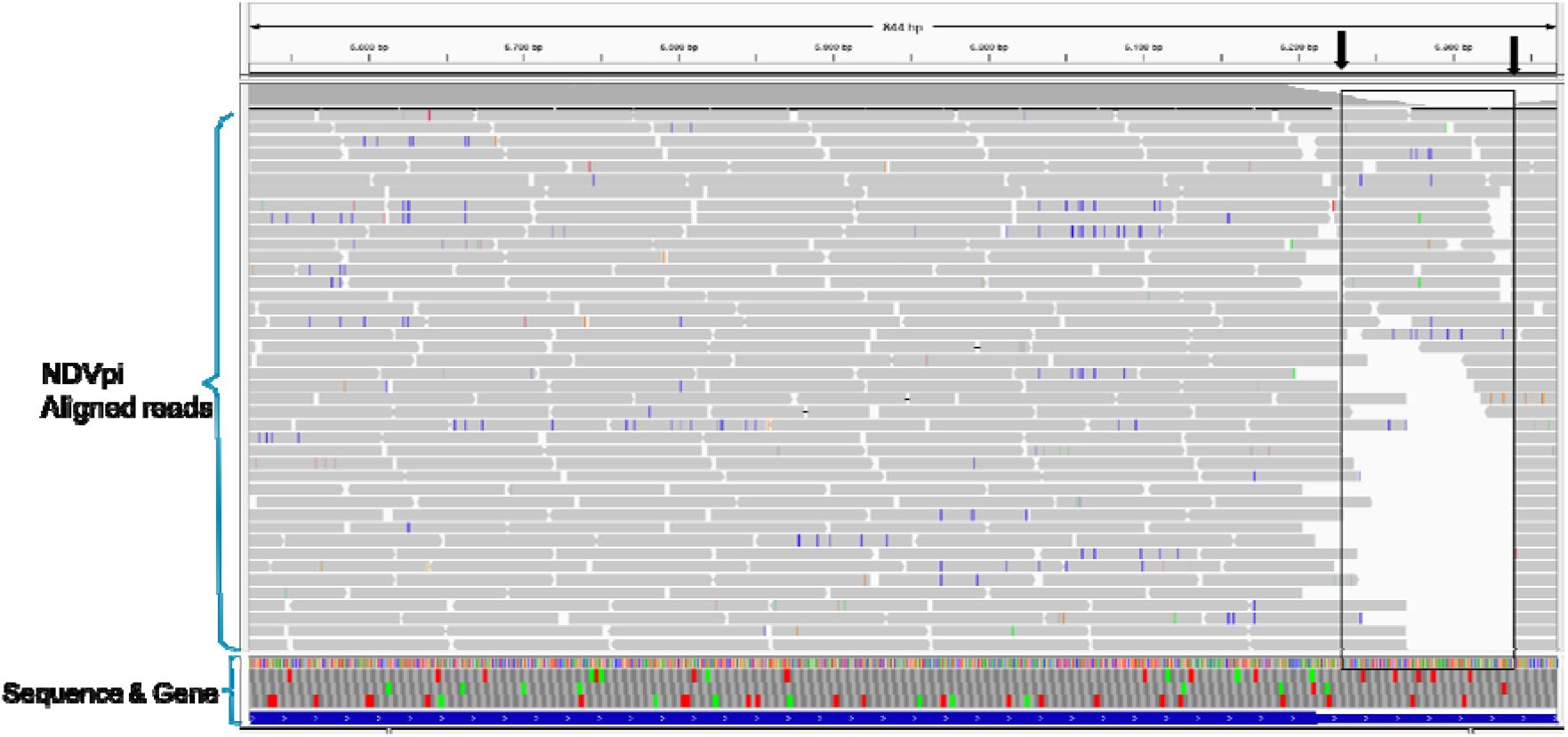
Alignment of NDVpi to the reference with truncated genome. Truncation of the viral genome is shown by the arrows. The truncated area was mapped to nucleotide position from 6203 to 6342 in L protein.

### Implications for NDV Persistence in Cancer Cells

The various genomic mutations and truncations identified in NDVpi provide insight into the molecular mechanisms that facilitate persistent infection in bladder cancer cells. The mutations in the NP, M, and P genes likely contribute to reduced replication efficiency and viral assembly, while the truncations in the HN and L genes may impair the virus’s ability to effectively attach to and replicate within host cells. This allows the virus to maintain a persistent, low-level infection, evading complete clearance by the host’s immune system. Similar mechanisms of persistence have been reported in other oncolytic viruses, such as vesicular stomatitis virus (VSV) and reovirus[10, 14].

Persistent infection poses a challenge for the use of NDV as an oncolytic virus in cancer therapy. While persistent infection may allow NDV to remain within the tumor microenvironment, it can reduce the cytolytic effects needed for tumor clearance. Previous studies have suggested that targeting the specific genomic regions involved in persistence could enhance NDV’s oncolytic potential[8]. The development of engineered NDV strains that minimize the likelihood of persistence, while maintaining robust cytolytic activity, represents a promising approach to overcoming this limitation[9].

## 4. Conclusion

This study identified key genomic alterations in the persistently infecting oncolytic Newcastle disease virus (NDVpi) in bladder cancer cells, specifically within the NP, M, P, HN, and L genes. These alterations include truncations, deletions, and insertions, suggesting that defects in these critical viral proteins may contribute to viral persistence by impairing viral replication, assembly, and host cell attachment. The mutations in the NP, M, and P genes, along with the truncations in the HN and L genes, appear to significantly affect the virus’s ability to effectively complete its replication cycle, potentially allowing the virus to evade immune detection and establish long-term persistence in the host cells.

However, the precise mechanisms by which these mutational changes lead to persistent infection remain unclear. Further investigation is needed to determine how these mutations arise and how they influence viral behavior, as well as to explore strategies to prevent persistence. Understanding these mechanisms is crucial for improving the therapeutic potential of NDV as an oncolytic agent in cancer therapy, enabling the development of targeted approaches to mitigate viral persistence and enhance treatment efficacy.

## Acknowledgments

None

## Author Contributions

A.V., U.A., C.S.C., C.D., S.A., and Y.K., designed the study. U.A. performed the wet and dry lab work as well as analysed the data. U.A wrote the paper. All authors reviewed the manuscript.

## Data Availability

All data supporting the results in this article are included in the present and supplementary files except the raw and processed sequencing data that have been submitted to the NCBI Sequence Read Archive with accession number: PRJNA543209 (https://www.ncbi.nlm.nih.gov/bioproject/PRJNA543209) and Gene Expression Omnibus (GEO) with accession number GSE140902 (https://www.ncbi.nlm.nih.gov/geo/query/acc.cgi?acc=GSE140902).

## Abbreviations

## Supplementary

**Supplementary 1.**
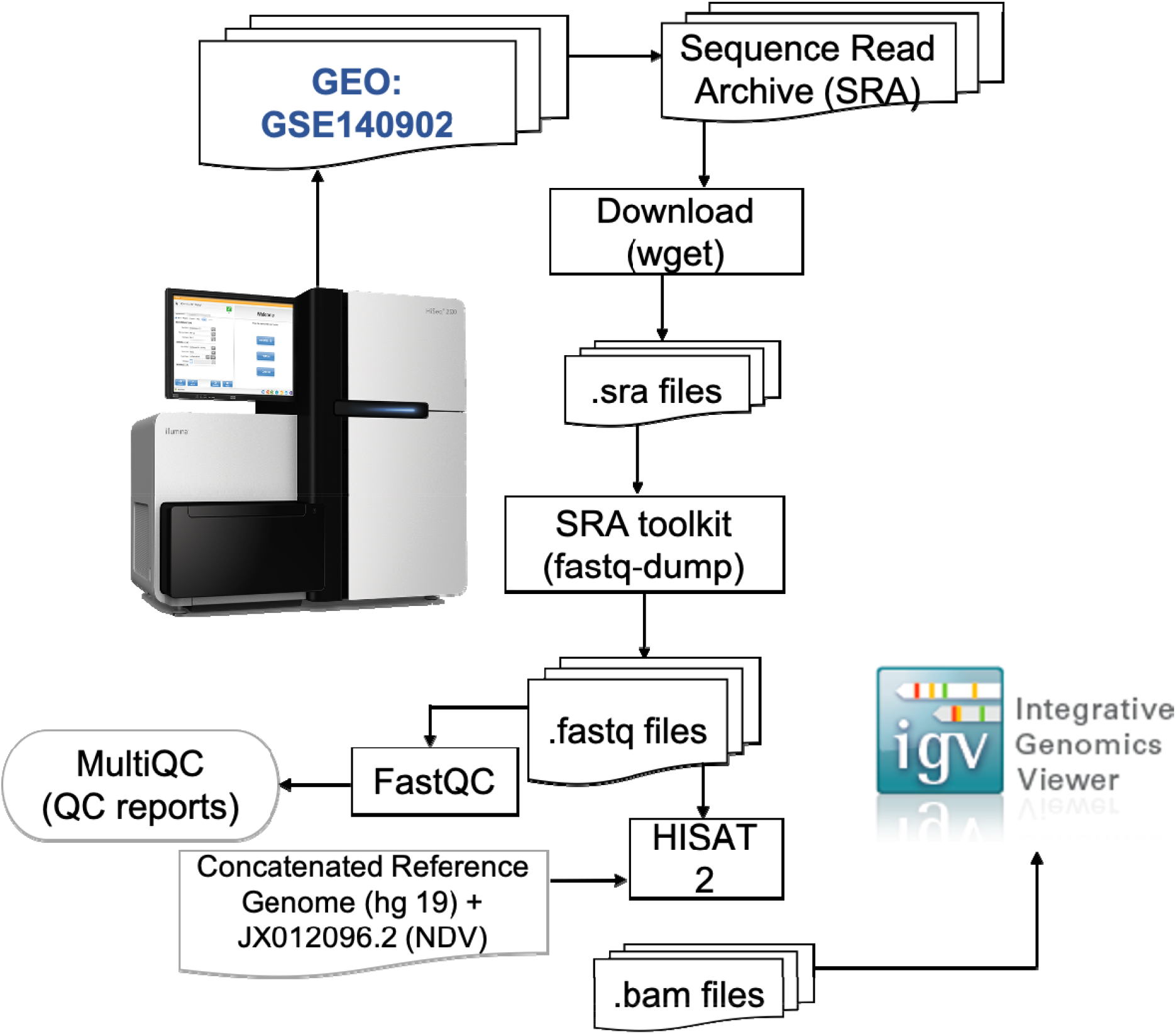
Workflow and bioinformatics pipelines used for the analysis

## References

1. Seal, B.S., D.J. King, and H.S. Sellers, The avian response to Newcastle disease virus. Developmental & Comparative Immunology, 2000. 24(2-3): p. 257–268.

2. Chellappa, M.M., et al., Complete genome sequence of Newcastle disease virus mesogenic vaccine strain R2B from India. 2012, Am Soc Microbiol.

3. de Leeuw, O. and B. Peeters, Complete nucleotide sequence of Newcastle disease virus: evidence for the existence of a new genus within the subfamily Paramyxovirinae. Journal of General Virology, 1999. 80(1): p. 131–136.

4. Phillips, R., A. Samson, and P. Emmerson, Nucleotide sequence of the 5′-terminus of Newcastle disease virus and assembly of the complete genomic sequence: agreement with the “rule of six”. Archives of virology, 1998. 143: p. 1993–2002.

5. Oberdorfer, A. and O. Werner, Newcastle disease virus: detection and characterization by PCR of recent German isolates differing in pathogenicity. Avian Pathology, 1998. 27(3): p. 237–243.

6. Kumar, R., et al., Velogenic newcastle disease virus as an oncolytic virotherapeutics: in vitro characterization. Applied biochemistry and biotechnology, 2012. 167: p. 2005–2022.

7. Shnyrova, A.V., et al., Vesicle formation by self-assembly of membrane-bound matrix proteins into a fluidlike budding domain. The Journal of cell biology, 2007. 179(4): p. 627–633.

8. Russell, S.J., K.-W. Peng, and J.C. Bell, Oncolytic virotherapy. Nature biotechnology, 2012. 30(7): p. 658–670.

9. Zamarin, D. and P. Palese, Oncolytic Newcastle disease virus for cancer therapy: old challenges and new directions. Future microbiology, 2012. 7(3): p. 347–367.

10. Chia, S.-L., K. Yusoff, and N. Shafee, Viral persistence in colorectal cancer cells infected by Newcastle disease virus. Virology journal, 2014. 11: p. 1–8.

11. Kelly, E.J., et al., Attenuation of vesicular stomatitis virus encephalitis through microRNA targeting. Journal of virology, 2010. 84(3): p. 1550–1562.

12. Doi, T., et al., Measles virus induces persistent infection by autoregulation of viral replication. Scientific reports, 2016. 6(1): p. 37163.

13. Swanton, C., Intratumor heterogeneity: evolution through space and time. Cancer research, 2012. 72(19): p. 4875–4882.

14. Goradel, N.H., et al., Oncolytic virotherapy: Challenges and solutions. Current problems in cancer, 2021. 45(1): p. 100639.

15. Msaouel, P., et al., Engineered measles virus as a novel oncolytic therapy against prostate cancer. The Prostate, 2009. 69(1): p. 82–91.

16. Vijayakumar, G., S. McCroskery, and P. Palese, Engineering Newcastle disease virus as an oncolytic vector for intratumoral delivery of immune checkpoint inhibitors and immunocytokines. Journal of Virology, 2020. 94(3): p. 10.1128/jvi.01677-19.

17. Andrews, S., FastQC: a quality control tool for high throughput sequence data http://www.bioinformatics.babraham.ac.uk/projects/fastqc. Babraham Bioinformatics, 2010.

18. Kim, D., B. Langmead, and S.L. Salzberg, HISAT: a fast spliced aligner with low memory requirements. Nature methods, 2015. 12(4): p. 357–360.

19. Robinson, J.T., et al., Integrative genomics viewer. Nature biotechnology, 2011. 29(1): p. 24–26.

20. Zhou, J., et al., Recent advancements in the diverse roles of polymerase-associated proteins in the replication and pathogenesis of Newcastle disease virus. Veterinary Research, 2025. 56(1): p. 8.

