## Supplementary figures and images for "Genomic alterations in persistently infecting oncolytic Newcastle disease virus reveal mechanisms of viral persistence in bladder cancer cells"

### Supplementary file

**Supplementary 1. Workflow and bioinformatics pipelines used for the analysis**

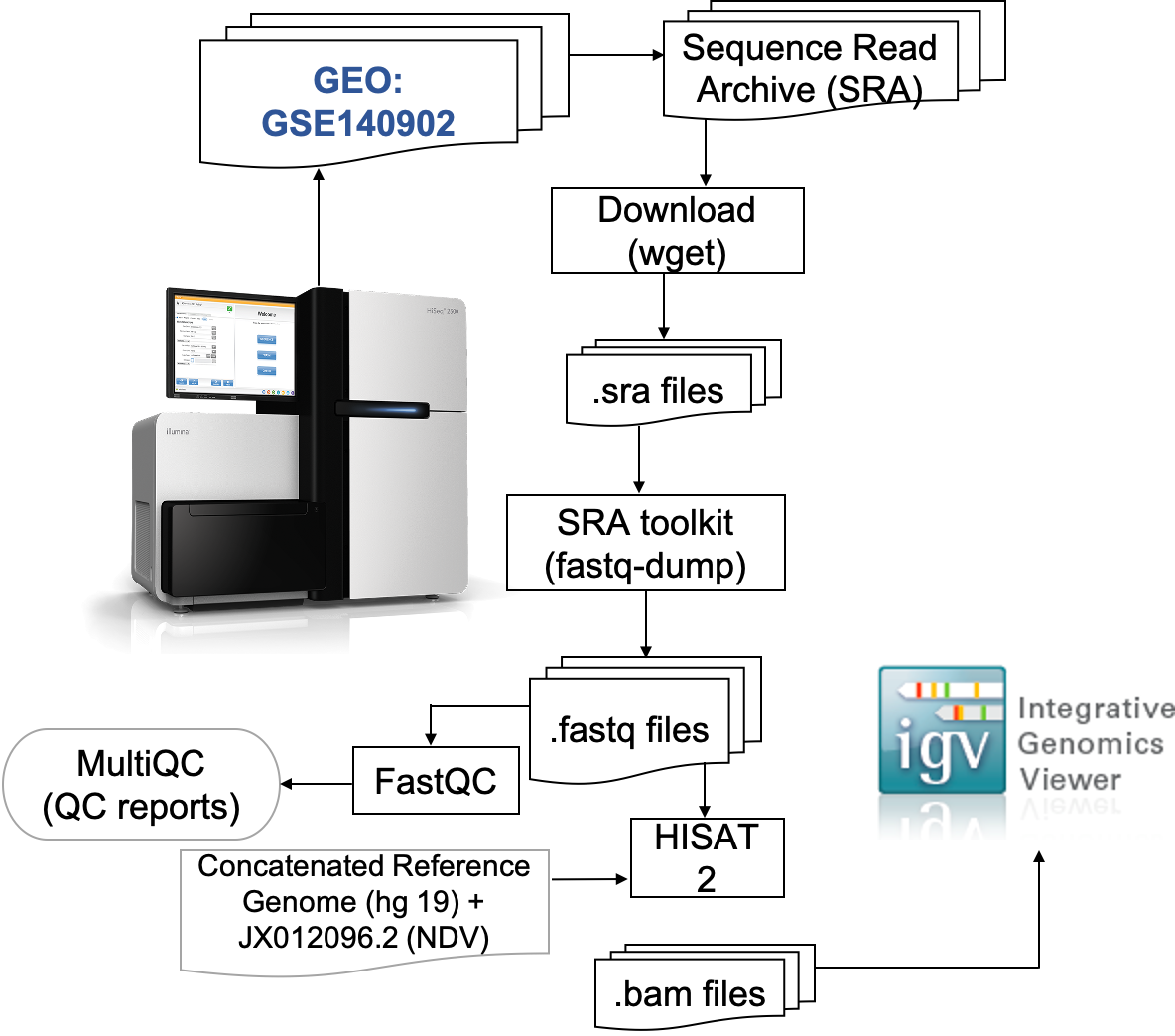
